# A secreted citrus protease cleaves an outer membrane protein of the Huanglongbing pathogen

**DOI:** 10.1101/2025.10.06.680679

**Authors:** Alexander J. McClelland, Bin Hu, Yuantao Xu, Chunxia Wang, Amelia H. Lovelace, Eva Hawara, Yuanchun Wang, Zhiqian Pang, Agustina De Francesco, Amit Levy, Nian Wang, Renier A. L. van der Hoorn, Qiang Xu, Wenbo Ma

**Affiliations:** The Sainsbury Laboratory, University of East Anglia, Norwich Research Park, Norwich, NR4 7UH, UK; National Key Laboratory for Germplasm Innovation & Utilization of Horticultural Crops, Huazhong Agricultural University, Wuhan, 430070, China; Citrus Research and Education Center, Department of Microbiology and Cell Science, University of Florida/IFAS, Lake Alfred, Florida, 33850, USA; Department of Microbiology and Plant Pathology, University of California Riverside, Riverside, CA 92521, USA; Plant Chemetics Laboratory, Department of Biology, University of Oxford, South Parks Road, Oxford, OX1 3RB, UK

**Keywords:** citrus greening disease, plant defense, plant innate immunity, plant-pathogen interactions, bacterial pathogens, Candidatus Liberibacter

## Abstract

Plants secrete a variety of proteases as a defense response during infection by microbial pathogens. However, the relationship between their catalytic activities and antimicrobial functions remains largely unknown. Particularly, few biologically relevant substrates of these proteases have been identified. Huanglongbing (HLB) has been a major threat to the citrus industry worldwide. The HLB-associated bacterium, Candidatus Liberibacter asiaticus (Las), was previously shown to deploy an inhibitor of papain-like cysteine proteases (PLCPs) to promote disease in citrus. In this study, we identified an outer membrane protein (OMP) of Las, LasOMP1, as a substrate of the citrus PLCP CsRD21a. LasOMP1 is one of the most highly expressed genes in Las. CsRD21a cleaves LasOMP1 and produces cleaved peptide products, which could be detected in vitro and in HLB-diseased citrus plants. We found that CsRD21a targets the N-terminal portion of LasOMP1, potentially at an extracellular loop region. Importantly, transgenic sweet orange overexpressing CsRD21a showed reduced Las titers and delayed disease symptoms, highlighting that engineering this protease is a promising strategy to enhance HLB resistance in citrus. Together, our work reveals a pathogen-derived substrate of plant PLCPs and highlights bacterial OMPs as direct targets of plant defense.

**Significance Statement:** Huanglongbing (HLB) is the most devastating disease of citrus with no resistance having been identified in commercial cultivars. Previous work implicated papain-like cysteine proteases (PLCPs) as an important hub of defense in citrus; however, their precise role in HLB tolerance remained unclear. Here, we identify and characterize an outer membrane protein (OMP) from the HLB-associated bacterium as a substrate of the citrus PLCP RD21a. We demonstrate a specific cleavage of the bacterial OMP by citrus RD21a, which may impair pathogen growth and/or activate plant immunity. Importantly, overexpression of RD21a enhances HLB tolerance in sweet oranges. This work identifies OMPs as substrates of plant PLCPs and provides insights into protease defense functions.

## Introduction

Plant hosts deploy a myriad of hydrolytic enzymes to suppress pathogen virulence. Among them, many proteases are secreted and have established functions in plant defense (1–3). For example, the papain-like cysteine protease (PLCP) RD21a from *Arabidopsis thaliana* and its homologs in rice and wheat contribute to defense against a range of pathogens, including nematodes, protists, viruses, and fungi (4). In line with this defense role, evolutionarily distant plant pathogens commonly deploy virulence proteins, known as effectors, that possess protease inhibitor activities as a counter-defense strategy (1). Furthermore, proteases can be co-opted by plant immune receptors to function as guarded decoys that trigger receptor-mediated defense upon inhibition by pathogen effectors (5). However, specific substrates of the proteases that directly contribute to their defense functions remain poorly understood.

Extracellular proteases can cleave both endogenous substrates and pathogen-derived substrates. Cleavage of endogenous substrates may release immunogenic damage-associated molecular patterns (DAMPs) or activate other proteases with defense functions (6, 7). On the other hand, cleavage of pathogen-derived substrates may directly lead to antimicrobial activities and/or release pathogen-associated molecular patterns (PAMPs) that activate cell surface receptor-mediated defense, called PAMP-triggered immunity (PTI). To date, only a few pathogen-derived substrates of secreted plant proteases have been identified. PC2, a small, secreted protein from the oomycete pathogen *Phytophthora infestans*, is cleaved by a host subtilase, releasing a peptide that triggers cell death and thus restricts pathogen growth (8). Soybean secretes an aspartic protease that degrades a *Phytophthora sojae* cell wall-degrading enzyme, XEG1, thus contributing to resistance (9). Arabidopsis aspartic proteases cleave a periplasmic protein, MucD, from the bacterial pathogen *Pseudomonas syringae*, thus directly inhibiting bacterial growth (10). Lastly, the plant subtilase SBT5.2 balances immune responses by both releasing and degrading PAMPs to activate and suppress defense, respectively (11, 12). The identification of protease substrates offers a unique opportunity to uncover mechanisms of plant immunity and provides important insights into engineering strategies for disease resistance in crops.

Huanglongbing (HLB), also known as citrus greening, is the most devastating citrus disease to which all commercial citrus varieties are susceptible (13, 14). HLB is primarily caused by gram-negative bacteria belonging to ‘*Candidatus* Liberibacter’ species, including *Ca.* Liberibacter asiaticus (Las), *Ca.* Liberibacter americanus (Lam), and *Ca.* Liberibacter africanus (Laf), with Las being the most widespread and impactful to the citrus industry. *Ca.* Liberibacter species are specialized, obligate bacteria transmitted by insect vectors, called psyllids, which deposit the bacteria into the plant phloem during feeding. In addition to the HLB-associating bacteria, *Ca.* L. solanacearum (Lso) causes disease in tomato, potato, pepper, and carrot (13, 15, 16). These bacteria have undergone significant genome loss compared to their pathogenic and non-pathogenic relatives in the Rhizobiaceae family, resulting in their inability to be cultured in vitro (17, 18). Each of these pathogens colonizes the plant phloem tissue, where mechanisms of plant defense responses are poorly understood.

Our previous characterization of Las effectors identified Sec-delivered effector 1 (SDE1) as a PLCP- inhibitor, implicating a role of PLCPs in phloem-based defense responses (19). SDE1 interacts with proteases from multiple PLCP subfamilies via their conserved protease domains, resulting in a decrease in their proteolytic activity in vitro and in planta (19). Transgenic citrus expressing SDE1 exhibited enhanced susceptibility to Las and accelerated HLB disease progression, indicating a correlation between PLCP inhibition and defense suppression (20). Furthermore, citrus PLCPs are induced during Las infection, consistent with their role as important nodes of defense in the phloem (19, 21, 22). Yet the mechanisms underlying PLCP defense functions are unclear.

In this work, we investigated the roles of PLCPs in citrus defense during Las infection. We found that *Citrus sinensis* RD21a (*Cs*RD21a) specifically interacts with an outer membrane protein (OMP) of Las named OMP1. This interaction leads to the cleavage of LasOMP1 near its N-terminal region. LasOMP1 is among the most highly expressed genes in Las and is also conserved across Liberibacter species, suggesting a critical role in bacterial survival and/or host colonization. Furthermore, LasOMP1 cleavage by PLCPs resulted in induced expression of a defense-related gene, leading to the possibility that an immunogenic peptide may be released to activate plant immunity. Finally, we generated transgenic citrus plants that overexpress *Cs*RD21a, which showed enhanced resistance to HLB as reflected by decreased Las titers and weakened disease symptoms. Together, these results shed light on the mechanisms of protease-based defense in an economically important disease and reveal bacterial OMPs as targets of plant immune system.

## Results

### Citrus PLCPs interact with a highly expressed outer membrane protein of Las

The exposed surface of the outer membrane (OM) of gram-negative bacteria is enriched in β-barrel outer membrane proteins (OMPs) and lipopolysaccharide (23). OMPs provide structural support to the bacterial cell and form channels for the passage of molecules in and out of the cell. OMPs contain an N-terminal signal peptide (SP) that facilitates their secretion into the periplasm followed by an OM-embedded transmembrane β-barrel domain which contains periplasmic and extracellular loops (24). As the only exposed proteins on the bacterial cell surface, we hypothesized OMPs may be targets of plant defense mechanisms. Using a hidden Markov model (HMM)-based approach (25, 26), we predicted six Las β-barrel OMPs from a total of 1,017 proteins encoded in the Las genome (strain psy62) and modeled their protein structures without the N-terminal SPs using AlphaFold 3 (**Fig. 1A**). We further used DeepTMHMM topology predictions (26) to determine the number of the transmembrane β-strands in each predicted OMP. Two 16- stranded β-barrel OMPs were annotated as “porin” or “BamA”, and one 26-stranded β-barrel OMP was annotated as “LptD” based on orthology to characterized OMPs in other bacteria. In addition, three eight-stranded β-barrel OMPs were identified and subsequently named OMP1, OMP2, and OMP3, respectively.

**Figure 1.**
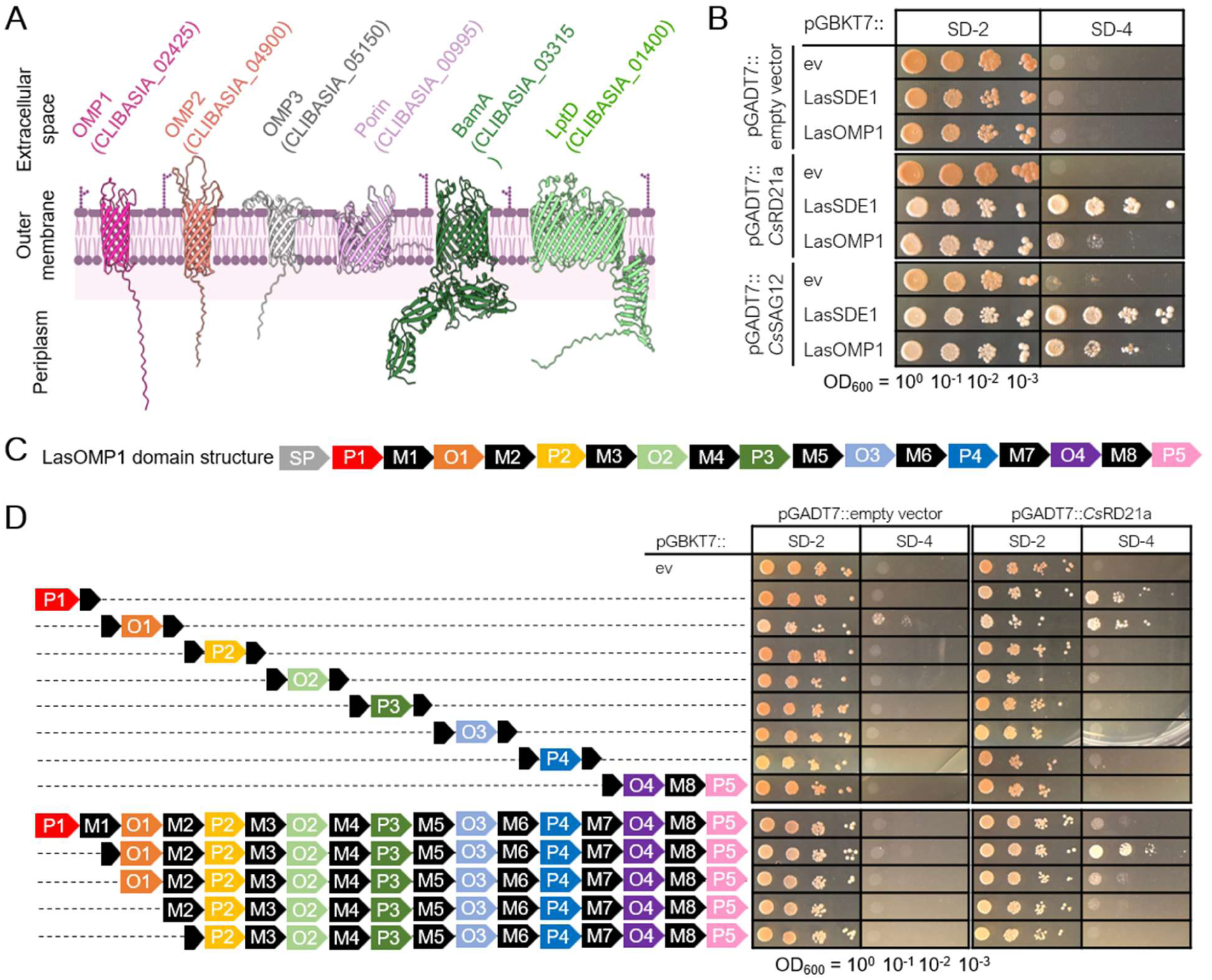
The citrus PLCP *Cs*RD21a interacts with the N-terminus of LasOMP1. (**A**) AlphaFold3 structural models of all Las OMPs without their signal peptides. All models were predicted with pTM scores between 0.65 and 0.84. The membrane image was sourced from BioRender.com. (**B**) Yeast-two-hybrid assay examining interactions between citrus PLCPs *Cs*RD21a and *Cs*SAG12 and Las membrane proteins. LasSDE1 and empty vectors (ev) served as positive and negative controls, respectively. SD-2 medium lacks leucine and tryptophan and was used to select the co-transformed colonies. One colony was serially diluted from OD_600_ = 1.0 and plated for each co-transformation in yeast on both SD-2 and SD-4, which lacks leucine, tryptophan, adenine, and histidine. Growth on SD-4 indicates protein-protein interaction. All proteins were expressed without their signal peptides. (**C**) LasOMP1 domain architecture (not to scale) predicted by OMPdb and DeepTMHMM. SP = signal peptide, P = periplasmic, M = membrane-bound, and O = outside/extracellular. (**D**) Yeast-two-hybrid assay examining interactions of *Cs*RD21a with LasOMP1 truncations. Dashed lines indicate deleted portions of LasOMP1, and the signal peptide of LasOMP1 was excluded.

To investigate potential interactions between citrus PLCPs and Las OMPs, we performed yeast two-hybrid assays using *Cs*RD21a and *Cs*SAG12 as the baits. We found that both *Cs*RD21a and *Cs*SAG12 interacted with OMP1, hereafter LasOMP1, but not with the other OMPs or an inner-membrane protein YajC (**Fig. 1B, S1A**). The relatively weaker interactions observed between the citrus PLCPs and LasOMP1 compared to LasSDE1 may indicate a less stable association, possibly due to proteolytic activity of the PLCPs against LasOMP1. Using publicly available transcriptomic data (27, 28), we found that LasOMP1 is the most highly expressed OMP and one of the most highly expressed Las genes overall in both grapefruit and psyllids (**Fig. S1B**). This expression profile suggests that LasOMP1 may have important biological functions in bacterial growth and host colonization.

Using OMPdb and DeepTMHMM (26, 29), we generated a LasOMP1 domain map (**Fig. 1C**). Following the N-terminal SP, LasOMP1 contains five periplasmic (P) domains, eight membrane-bound (M) domains, and four extracellular/outside (O) domains. Using this domain architecture as a guide, we generated eight truncations, dividing LasOMP1 at each of the first seven membrane-bound domains (M1-M7). The truncations were then tested for interaction with *Cs*RD21a via yeast-two-hybrid. Two truncations near the N-terminus of LasOMP1, P1 and O1, were still able to interact with *Cs*RD21a, although the O1 truncate exhibited a weak autoactivation (**Fig. 1D**). To further confirm *Cs*RD21a association with the N-terminal region of LasOMP1, a series of truncated mutants were generated and tested for interactions with *Cs*RD21a. The results showed that the interaction with *Cs*RD21a was abolished only when both P1 and O1 domains were removed (**Fig. 1D**), demonstrating that the N-terminal P1 and OI regions of LasOMP1 are both sufficient and necessary for interaction with host PLCP *Cs*RD21a.

### *Cs*RD21a cleaves LasOMP1 in a semi-in vitro assay

To assess whether LasOMP1 is a substrate of *Cs*RD21a, we established a semi-in vitro protease cleavage assay (**Fig. 2A**). It has been previously demonstrated that Arabidopsis RD21a undergoes both N- and C-terminal processing during its activation and secretion (30, 31). Therefore, we opted to express *Cs*RD21a without a tag and confirmed its enzymatic activity using activity-based protein profiling (ABPP). As a negative control, a catalytic mutant of *Cs*RD21a, *Cs*RD21a^CHN^, was generated in which each of its three catalytic residues (C163, H299, and N319) were replaced with alanine. Full-length *Cs*RD21a and *Cs*RD21a^CHN^ were expressed via *Agrobacterium*-mediated transient expression in *Nicotiana benthamiana,* and their secreted proteins (lacking an SP) were collected from apoplastic fluid (AF) and incubated with the biotinylated ABPP cysteine protease probe DCG-04 (32). Using streptavidin-HRP, we detected clear DCG- 04 labeling signals of *Cs*RD21a between 25 and 35 kDa in leaves expressing the wild-type protein (**Fig. 2B**), corresponding to active proteoforms that have undergone removal of the autoinhibitory prodomain (**Fig. S2**). This signal decreased in the presence of the cysteine protease inhibitor E-64 in a dosage-dependent manner, confirming that the enzymatic activity detected was indeed from *Cs*RD21a (**Fig. 2B**). As expected, no ABPP signal was observed in samples expressing *Cs*RD21a^CHN^.

**Figure 2.**
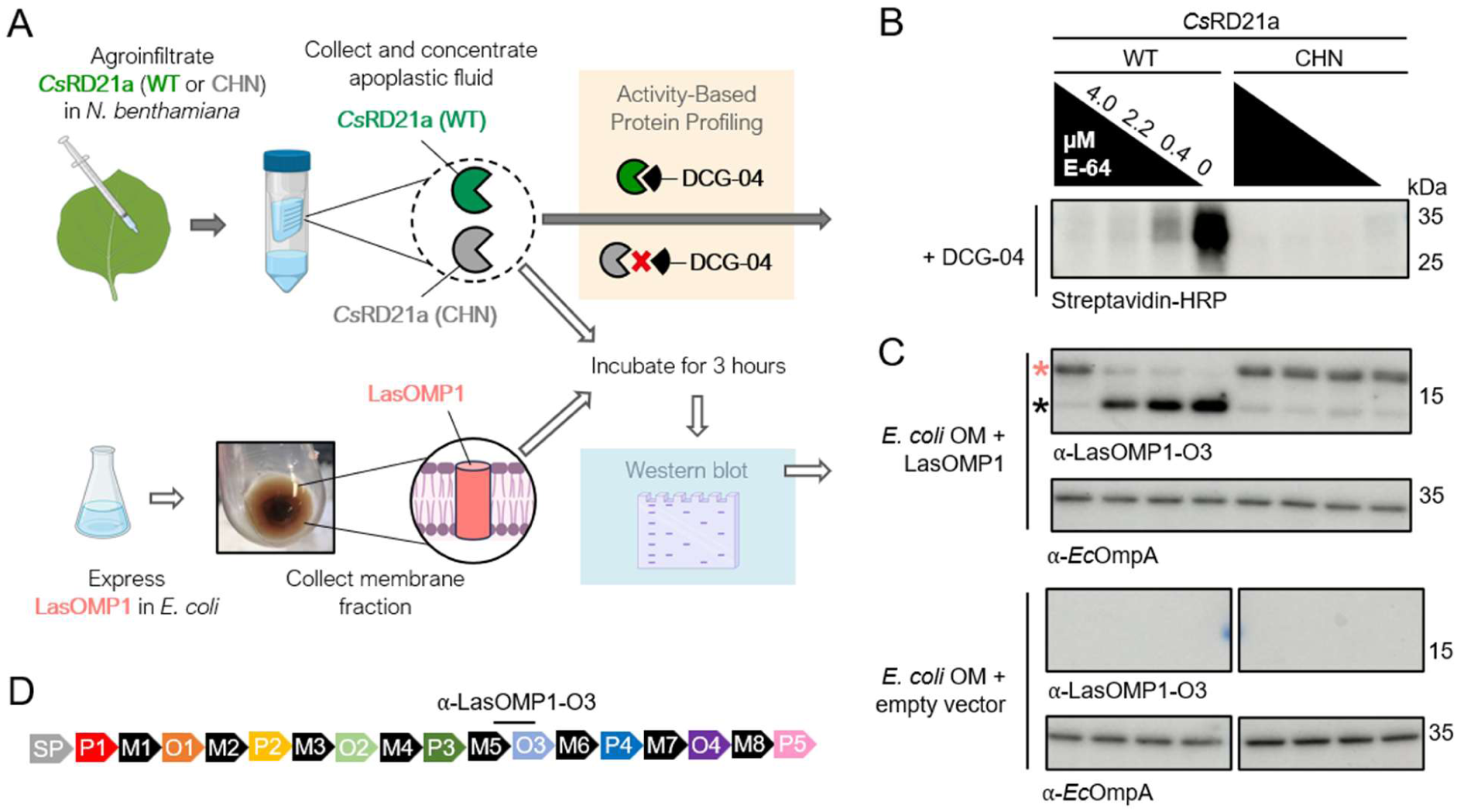
*Cs*RD21a cleaves LasOMP1 in a semi-in vitro assay. (**A**) Workflow of the semi-in vitro and activity-based protein profiling (ABPP) assays made using BioRender.com. (**B**) Western blot detection of wild-type (WT) or catalytically dead (CHN) *Cs*RD21a in apoplastic fluid (AF) using ABPP, where a concentration gradient of the cysteine protease inhibitor E-64 was incubated with the AF prior to DCG-04 labeling. Western blot detection of DCG-04-bound proteases was performed using Streptavidin-HRP. (**C**) Cleavage of LasOMP1 by *Cs*RD21a. E-64- or DMSO-treated AF containing *Cs*RD21a WT or the CHN mutant was incubated with *E. coli* membrane fractions containing LasOMP1. *Ec*OmpA was used as a loading control. Both OMPs were detected by Western blotting using protein-specific antibodies. The band representing full-length LasOMP1 is marked with a pink asterisk, and cleaved product is marked with a black asterisk. Membrane fractions extracted from *E. coli* carrying the empty vector were used as the negative control. (**D**) Domain structure of LasOMP1 showing the binding site of the α-LasOMP1-O3 antibody.

To test if LasOMP1 can be cleaved by *Cs*RD21a, the AF extracts containing the active or catalytic mutant of *Cs*RD21a were incubated with LasOMP1 produced in *E. coli* (**Fig. 2A**). To maximize LasOMP1 secretion, we replaced its native SP with the *E. coli* OmpA SP. Membrane fractions of *E. coli* were collected from LasOMP1-expressing cells using cells carrying the empty vector as a control. *Ec*OmpA was detected with an *Ec*OmpA-specific antibody, confirming the enrichment of bacterial OMPs in the samples (**Fig. 2C**). LasOMP1 was detected using a LasOMP1-specific antibody (⍺-LasOMP1-O3), which was raised against the third extracellular loop (O3) (**Fig. 2D**). These bacterial membrane protein extracts were incubated with *Cs*RD21a-containing AF for three hours at room temperature. LasOMP1 appeared in its mature, secreted form (lacking the SP) (∼19.2 kDa) when incubated with wild-type *Cs*RD21a and a high concentration of E- 64 (4.0 µM) (**Fig. 2C**). However, a lower band below 15 kDa appeared, with a significant reduction of full-length LasOMP1 protein, in the presence of low concentrations of E-64. E-64 did not affect LasOMP1 stability when incubated with *Cs*RD21a^CHN^, suggesting that LasOMP1 is proteolytically cleaved only by wild-type *Cs*RD21a. Unlike LasOMP1, *Ec*OmpA was not affected by *Cs*RD21a (**Fig. 2C**), further demonstrating that the cleavage of LasOMP1 by *Cs*RD21a was specific.

### *Cs*RD21a interacts with OMP1 homologs in Liberibacter

To further understand LasOMP1 as a putative target of plant defense, we identified its homologs in other Liberibacter species, which include the citrus pathogens Laf and Lam, the potato pathogen Lso, and a non-pathogenic, culturable relative, *L. crescens* (Lcr) (**Table S1**). Phylogenetic analysis of 31 HMM-predicted Liberibacter OMPs revealed conserved clades corresponding to LptD, Porin, and BamA, which each contain one protein from each species (**Fig. 3A**). Interestingly, we observed a conserved and expanded cluster comprising LasOMP1 and a single representative homolog from each Liberibacter proteome, apart from Lcr which has four homologs (**Fig. 3A, B**). In contrast, OMP2 and OMP3 are specific to pathogenic *Ca.* Liberibacter spp., suggesting they may have a role in pathogenesis.

**Figure 3.**
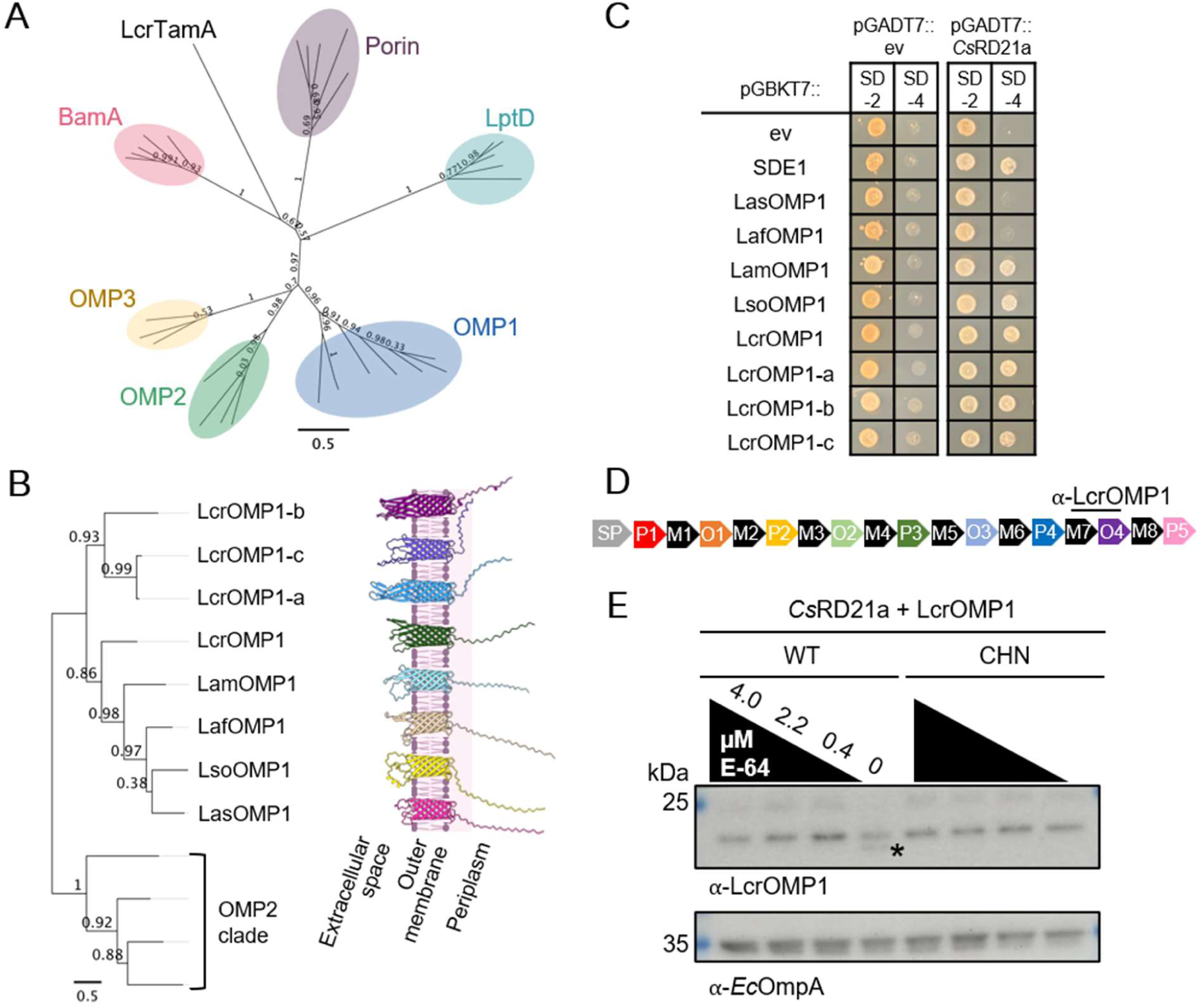
*Cs*RD21a interacts with OMP1 homologs in *Liberibacter* spp. and weakly cleaves LcrOMP1. (**A**) Unrooted maximum likelihood tree depicting six clusters of 31 predicted OMPs in Liberibacter (Table S1). A MAFFT amino acid sequence alignment was used to generate the phylogeny. FastTree support values are indicated at each node. Tree is visualized using Geneious Prime. (**B**) Phylogenetic tree produced from a MAFFT alignment of Liberibacter OMP1 proteins, with the OMP2 cluster as an outgroup. AlphaFold3 structural models of the OMP1 proteins are shown beside the tree. All pTM scores for the models were between 0.65 and 0.73, and the membrane image was sourced from BioRender.com. (**C**) Yeast two-hybrid assays detecting interactions between OMP1 homologs with *Cs*RD21a. One colony was diluted to OD_600_ = 0.4 and plated for each co-transformation in yeast on both SD-2 and SD-4 media. All proteins were cloned without their signal peptides. (**D**) Domain structure of LcrOMP1 depicting the binding site of the α-LcrOMP1 antibody. (**E**) Semi-in vitro cleavage assay of LcrOMP1 by *Cs*RD21a. Wild-type (WT) and catalytic mutant (CHN) *Cs*RD21a were incubated with *E. coli* membrane fractions containing LcrOMP1. Cleaved product of LcrOMP1 is marked with the asterisk. *Ec*OmpA was used as a loading control.

In Las, *OMP1* is one of the most highly expressed genes in both the plant host and the insect vector (**Fig. S1B**). To determine if OMP1 is also the dominant OMP in Lcr, we performed RNAseq analyses of this bacterium grown in liquid medium. This analysis revealed that LcrOMP1 homologs are also the highest expressed OMPs and each LcrOMP1 paralog was found in the top 17% of the most abundant Lcr transcripts (**Fig. S3**). The LcrOMP1 homologs form a similar AlphaFold-predicted protein model in which the transmembrane β-barrel and periplasmic disordered region are structurally conserved (**Fig. 3B**). However, their extracellular loops appear to have diverged and, in some cases, expanded. Indeed, sequence alignment of amino acid sequences clearly shows a lower degree of conservation in the O domains as compared to M domains, indicating potential functional diversification (**Fig. S4**).

The expression pattern, conservation, and expansion of OMP1 proteins suggests that they may play an important role in bacterial growth and thus could serve as targets of plant defense. These observations prompted us to further study this family of proteins as putative *Cs*RD21a substrates. We found that *Cs*RD21a interacts with each tested OMP1 homolog using yeast-two-hybrid assays (**Fig. 3C**). Both LasOMP1 and LafOMP1 exhibit weak interactions with *Cs*RD21a, as evidenced by less yeast growth on the selective media, compared to the other OMP1 proteins, which may indicate protease-substrate interactions.

To assess whether *Cs*RD21a cleaves the OMP1 homologs, we systematically tested them using the semi-in vitro assay. Because adding a C-terminal tag to OMPs for expression in *E. coli* obscured the identification of mature proteins from the membrane extracts, we generated an antibody, ⍺-LcrOMP1, based on the M7- O4 region of LcrOMP1 that would allow us to monitor potential cleavage of LcrOMP1 proteins (**Fig. 3D**). Incubation of AF containing wild-type *Cs*RD21a but not *Cs*RD21a^CHN^ with LcrOMP1 resulted in the appearance of a cleaved product only in the absence of the protease inhibitor E-64 (**Fig. 3E**). This is in stark difference with what we observed in LasOMP1, which was efficiently cleaved by *Cs*RD21a even in the presence of E-64 at a concentration as high as 2.2 µM. These results suggest that OMP1 proteins may be common targets of plant PLCPs, but the strength of *Cs*RD21a interaction, and the efficacy of cleavage, varies for different OMP1 homologs.

### *Cs*RD21a cleaves LasOMP1 at its N-terminus

Our yeast-two-hybrid results suggested that the N-terminal region of LasOMP1 mediates interaction with *Cs*RD21a. This region includes the O1 loop, which is predicted to be exposed to the cell surface of Las. Furthermore, cleavage in this region would produce a ∼14 kDa product, which is similar to what was detected using the ⍺-LasOMP1-O3 antibody. Therefore, we hypothesized that the O1 loop is cleaved by *Cs*RD21a. To test this, we synthesized a quenched octopeptide including one amino acid of the M1 domain and the following seven amino acids of the O1 loop of LasOMP1 (q-OMP1-O1) (**Fig. 4A**). Upon cleavage of the peptide, the N-terminal quencher would be separated from the C-terminal fluorophore, causing fluorescence (11, 12). We found that incubation of the wild-type *Cs*RD21a-containing AF with q-OMP1-O1 resulted in a significantly higher level of fluorescence compared to the *Cs*RD21a^CHN^ mutant (**Fig. 4B**). As a control, a quenched octopeptide derived from the bacterial flagellin flg22 (QP2) that was previously found to be processed by subtilases but not PLCPs (11) did not emit fluorescence that was different between wild-type *Cs*RD21a and *Cs*RD21a^CHN^ treatments. This result suggests that the O1 loop of LasOMP1 can be cleaved by *Cs*RD21a. We also examined the OMP1 homolog in the Las strain Ishi-1, which contains a single nucleotide polymorphism, resulting in a histidine to aspartic acid mutation at the 73^rd^ amino acid in the O1 loop (**Fig. S5A**). LasOMP1^Ishi-1^ could also be cleaved by *Cs*RD21a (**Fig. S5B**), suggesting that *Cs*RD21a is effective in targeting naturally occurring variants of OMP1 in Las.

**Figure 4.**
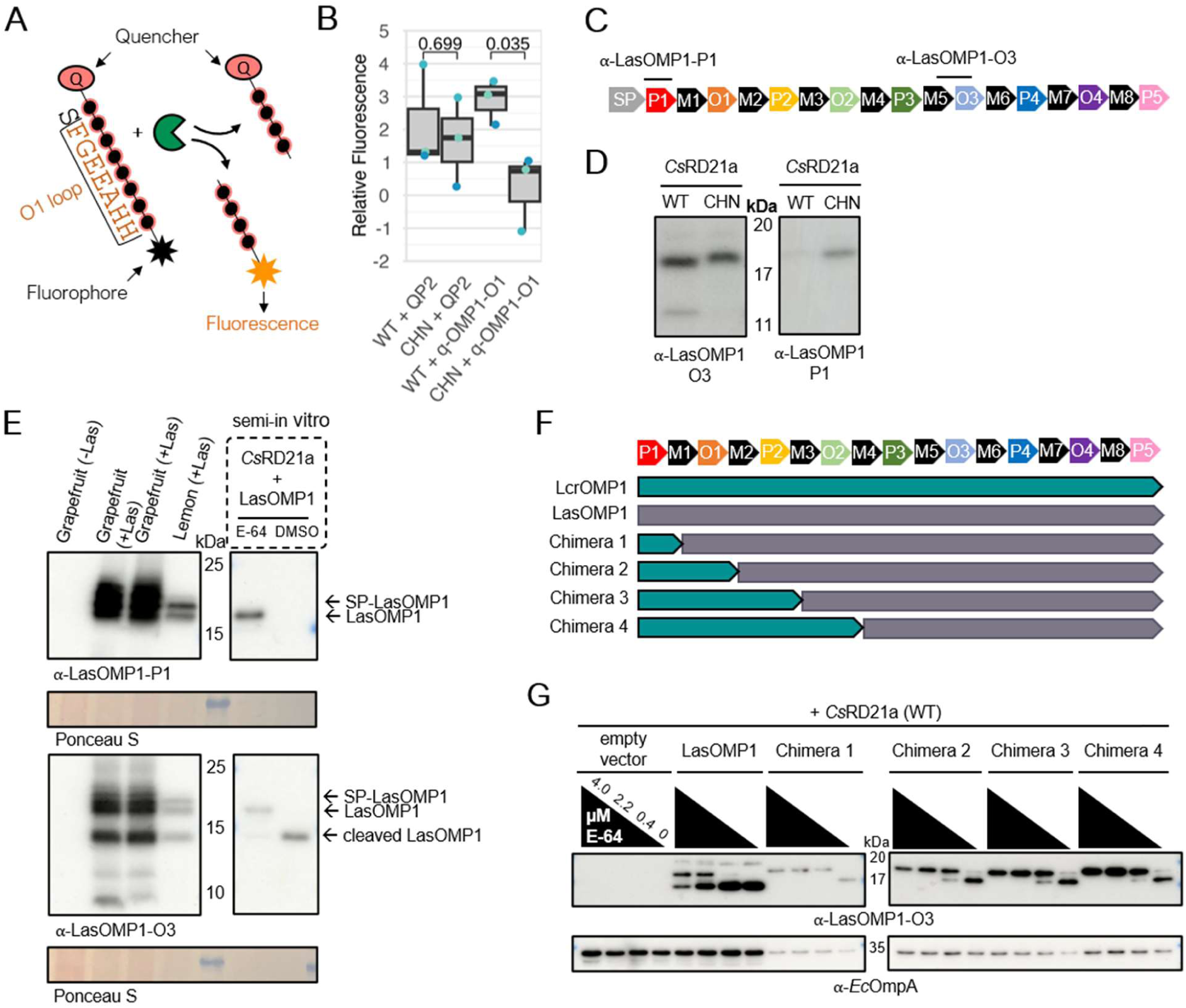
*Cs*RD21a cleaves within the N-terminal region of LasOMP1. (**A**) Experimental design of using a quenched octopeptide (qPeptide) to detect protease-based on cleavage. (**B**) *Cs*RD21a cleaves the O1 loop of LasOMP1. Apoplastic fluid containing wild-type (WT) or the catalytic mutant (CHN) of *Cs*RD21a was extracted from *N. benthamiana* and incubated with qPeptides corresponding to flg22 (QP2) or the O1 loop of LasOMP1 (q-OMP1-O1). Relative fluorescence unit (RFU) values from inhibited (E-64-treated) apoplastic fluid were subtracted from uninhibited (DMSO-treated) for each protease-qPeptide combination after 30 minutes, and the resulting values were normalized to the WT and CHN controls, respectively, without the qPeptides. Data from three independent biological replicates were presented (corresponding to the different colored data points), and the p-values shown between WT and CHN treatments were derived from two-tailed t-tests. (**C**) Schematic depicting the domain structure of LasOMP1 and the binding site of the α-LasOMP1-P1 antibody. (**D**) Western blots showing the different *Cs*RD21a wild-type (WT)- and catalytic mutant (CHN)-produced cleavage products detected by different LasOMP1-specific antibodies. (**E**) Western blotting of healthy (-Las) and infected (+Las) citrus seed coat vasculatures detecting LasOMP1. Samples were run alongside *Cs*RD21a-cleaved LasOMP1 from the semi-in vitro assay. Infected grapefruit samples are in duplicates. Ponceau S was used as the loading control. (F) Schematic depicting the LcrOMP1 and LasOMP1 domain structure and the chimeras used in semi-in vitro cleavage assays shown in panel G. (**G**) Western blots showing the cleavage of OMP1 chimeras by *Cs*RD21a detected by the α-LasOMP1-O3 antibody. *Ec*OmpA was used as a loading control.

We further characterized the cleavage of LasOMP1 by *Cs*RD21a using Western blotting. For this purpose, we generated an antibody against the P1 domain of LasOMP1, which should be in the proximity of the potential cleavage site (**Fig. 4C**). Notably, we were unable to detect any *Cs*RD21a-produced cleavage products of LasOMP1 using the ⍺-LasOMP1-P1 antibody, although the cleavage products were clearly detectable using the ⍺-LasOMP1-O3 antibody (**Fig. 4D**). Nonetheless, full-length LasOMP1 proteins were largely reduced in Western blots using the ⍺-LasOMP1-P1 antibody in the presence of wild-type *Cs*RD21a, suggesting that the cleavage likely occurs in the proximity of the antibody-binding site.

We next tested whether LasOMP1 is cleaved in citrus during Las infection using the ⍺-LasOMP1-O3 and ⍺-LasOMP1-P1 antibodies by Western blotting. Total proteins were extracted from HLB-infected citrus seed coat vasculatures, which had high titers of Las (33, 34). Two bands larger than 15 kDa appeared using both antibodies, likely corresponding to the SP-containing, non-secreted LasOMP1 protein (21.5 kDa) and the SP-lacking, secreted LasOMP1 protein (19.2 kDa), respectively (**Fig. 4E**). The 19.2 kDa band is the same size as the membrane-incorporated LasOMP1 proteins produced in *E. coli*. Interestingly, an additional <15 kDa band, likely representing the cleavage product by *Cs*RD21a, was detectable using the ⍺-LasOMP1- O3 antibody but not the ⍺-LasOMP1-P1 antibody, similar to what was observed in the semi-in vitro assay (**Fig. 4E**). No LasOMP1 signal was detectable in healthy citrus samples using these antibodies, confirming their specificity. Together, these results suggest that *Cs*RD21a cleaves LasOMP1 *in planta* during natural infection.

The relatively weaker cleavage of LcrOMP1 than LasOMP1 by *Cs*RD21a allowed us to further dissect the cleavage site of LasOMP1 by generating chimeras. We generated four chimeras that contained increasingly larger N-terminal fragments of LcrOMP1 and shorter fragments of the C-terminus of LasOMP1 while retaining the binding site of the ⍺-LasOMP1-O3 antibody (**Fig. 4F**). These chimeras were individually expressed in *E. coli* and subjected to the semi-in vitro cleavage assay. Intriguingly, all the chimeras exhibited reduced cleavage efficiency compared to LasOMP1 (**Fig. 4G**), indicating the N-terminal region of LcrOMP1, containing only the P1 domain and two amino acids of the adjacent M domain, is sufficient to reduce the cleavage efficiency by *Cs*RD21a. Together with the q-OMP1-O1 cleavage assay, our results suggest that LasOMP1 may cleave multiple sites in the N-terminal region, which is consistent with our previous finding that the P1 and O1 domains both mediate *Cs*RD21a interaction.

### *Cs*RD21a overexpression enhances Las tolerance in citrus

Upon HLB infection, proteases are dynamically regulated in citrus (21). PLCP expression is induced in the tolerant variety Sugar Belle mandarin but decreased in the susceptible variety Pineapple sweet orange (22), suggesting that PLCPs may contribute to HLB tolerance. We overexpressed *CsRD21a* without a tag (given its proteolytic processing *in planta*) under the 35S promoter in sweet orange. Increased transcript abundance of *CsRD21a* in the transgenic plants compared to wild-type plants was confirmed by RT-qPCR (**Fig. S6A**). Six independent transgenic lines with ∼20-fold higher expression of *CsRD21a* were examined in Las infection experiments. No visible differences in growth and development were observed between *CsRD21a*-overexpressing (*CsRD21a-*OE) and wild-type (WT) citrus plants under normal growth conditions. We then examined HLB tolerance of *CsRD21a*-OE and WT plants by grafting transgenic buds onto the same branches of HLB-infected sweet orange plants (**Fig. S6B**). Las titers were monitored monthly by qPCR in the scions. Las was first detected (CT < 35) in WT scions at 6 months post inoculation (mpi) but was delayed until 9 mpi in the *CsRD21a*-OE scions. In addition, the Las titers were significantly lower in *CsRD21a*-OE scions compared to WT (**Fig. 5A**). *CsRD21a*-OE scions also appeared to be healthier than WT scions over the monitoring period (up to 10 mpi) (**Fig. 5B**).

**Figure 5.**
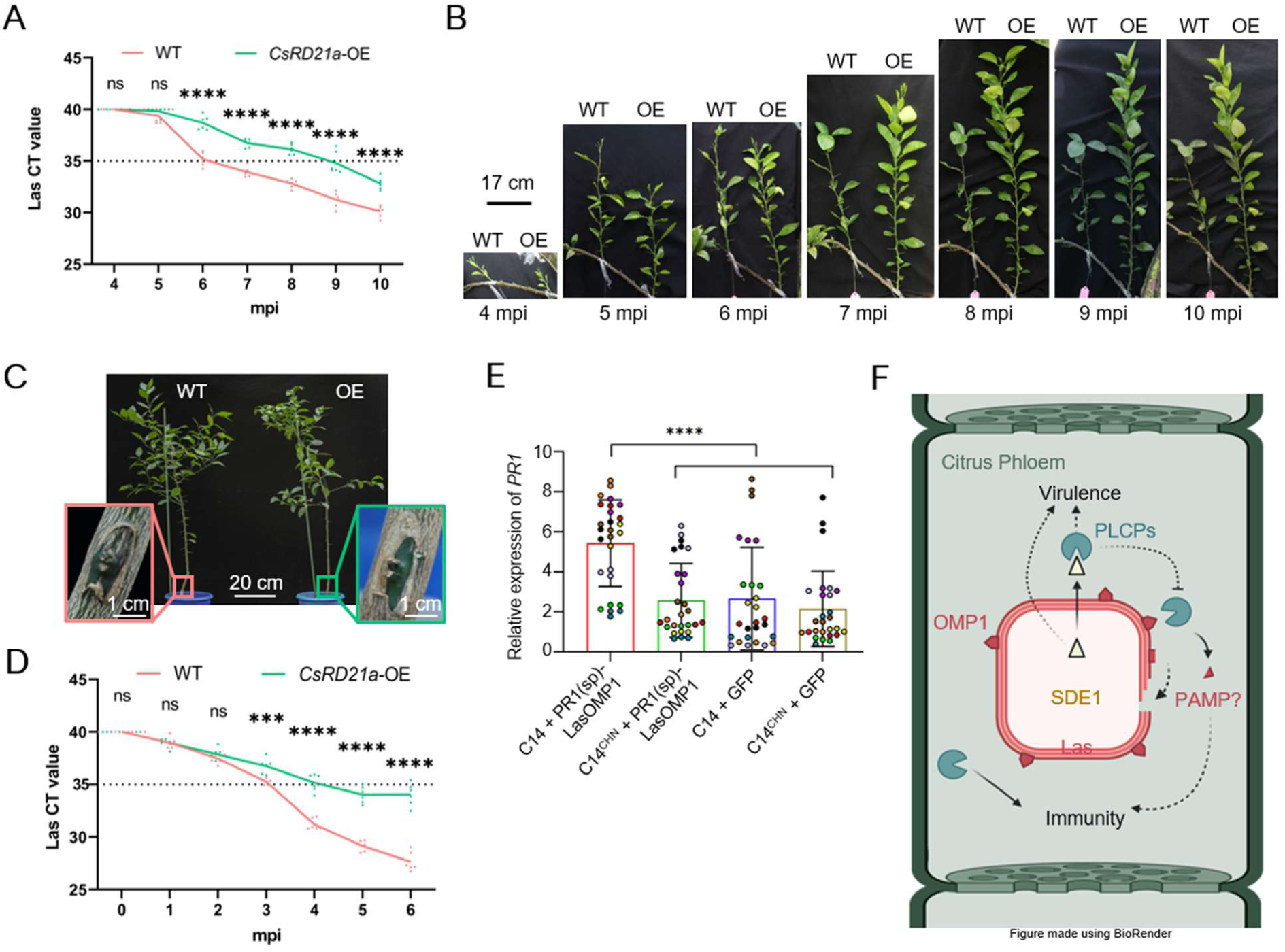
RD21a reduces Las growth in citrus and activates a LasOMP1-dependent immune response. (**A**) qPCR determination of Las titers in wild-type (WT) and transgenic sweet oranges overexpressing *CsRD21a* (OE), which were grafted on Las-infected citrus plants. Data points represent six independent biological replicates. Asterisks indicate significant differences: ns = no significance and **** = p < 0.0001. Las titer was determined at 4-10 months post inoculation (mpi). (**B**) Images of grafted WT or *CsRD21a*-OE scions on HLB-infected citrus. (**C**) Images of WT and *CsRD21a*-OE sweet orange plants graft-inoculated with Las-infected buds at nine months post inoculation. Pink and green boxes show the sites of graft-inoculation on WT and *CsRD21a*-OE plants, respectively. (**D**) qPCR determination of Las titers in WT and *CsRD21a*-OE sweet orange plants graft-inoculated with Las-infected buds. Data points represent six independent biological replicates. Asterisks indicate significant differences based on Student’s t-test: ns = no significance, *** = p < 0.001, and **** = p < 0.0001. (**E**) Relative expression of *PR1* determined by RT-qPCR in *N. benthamiana* leaves transiently co-expressing C14 (WT or CHN) and PR1(sp)-LasOMP1 or GFP. Nine independent biological replicates (shown by different colored data points) were analyzed. Three technical replicates were included for each biological replicate. Relative expression levels were calculated using the 2^−ΔΔCT^ method with *Actin* as the internal control. Data are presented as the mean ± standard deviation. Statistical significance was determined by one-way ANOVA, where **** = p < 0.0001. (**F**) Model of PLCP-dependent defense in the citrus during Las infection. PLCPs, such as *Cs*RD21a, contribute to HLB tolerance by targeting LasOMP1 for cleavage, which could directly inhibit Las growth. In addition, LasOMP1 cleavage leads to immune activation, potentially through the release of a PAMP. As a counter-defense, Las secretes an effector, SDE1, which acts as a protease inhibitor to block PLCP-dependent defense and contribute to pathogen virulence. Made using BioRender.com.

To further confirm the role of *CsRD21a* in defense during Las infection, we graft-inoculated *CsRD21a*-OE and WT citrus plants with Las-infected buds (**Fig. 5C**) and monitored Las titers over a 6-month period. In this assay, Las was first detected (CT < 35) in WT scions at 3 mpi but delayed to 4 mpi in *CsRD21a*-OE scions. In the time points when Las was detectable in both genotypes, their titers were significantly lower in *CsRD21a*-OE than WT citrus (**Fig. 5D**). Taken together, these results demonstrated that over-expression of *CsRD21a* reduced Las infection and HLB progression in citrus.

### RD21a-dependent cleavage of LasOMP1 may activate defense in *N. benthamiana*

Cleavage of LasOMP1 may contribute to defense through multiple mechanisms. Given the high expression level of *LasOMP1*, it may have housekeeping functions, thus its cleavage likely inhibits Las growth. Unfortunately, this cannot be directly tested since Las is unculturable. Furthermore, LasOMP1 cleavage may release peptide(s) that activates plant immunity. To test this, we co-expressed LasOMP1 in *N. benthamiana* with *Cs*RD21a or its tomato homolog C14. To ensure secretion of LasOMP1 into the apoplast, we replaced its SP with the pathogenesis-related protein 1 (PR1) SP. In this experiment, we consistently observed more efficient LasOMP1 cleavage by C14 than *Cs*RD21a (**Fig. S7A**). Indeed, C14, but not C14^CHN^, is highly active in *N. benthamiana* apoplastic fluid and cleaves LasOMP1 in the semi-in vitro assay (**Fig. S7B, C**). Therefore, we used C14 to test for potential immune activation upon LasOMP1 cleavage, using *PR1* expression as a proxy. C14 or C14^CHN^ did not induce *PR1* expression when co-expressed with GFP, indicating that its proteolytic activity alone could not activate immunity in *N. benthamiana* (**Fig. 5E, Fig. S7D**). In contrast, wild-type C14 co-expressed with LasOMP1 significantly induced *PR1* expression compared to C14^CHN^, suggesting that LasOMP1 can activate defense only when proteolytically processed. These findings suggest a possible mechanism in which RD21a could contribute to HLB tolerance by activating an immune response upon cleavage of LasOMP1 (**Fig. 5F**).

## Discussion

The class of RD21-like PLCPs has been implicated in plant immunity through genetic studies and the identification of numerous pathogen-derived inhibitors; yet, no biologically relevant substrates of these proteases have been previously identified (4). Indeed, no bacteria-derived substrates of any plant PLCP have yet been identified. Here, we identified bacterial OMPs as direct targets of plant PLCPs. Specifically, we observed that OMP1 from the citrus HLB pathogen Las is specifically cleaved by the citrus RD21a. LasOMP1 is one of the most highly expressed Las genes and is significantly more highly expressed than other Las OMPs. Similarly, our RNAseq analyses on the culturable relative of Las, Lcr, also revealed high expression of LcrOMP1, suggesting that this family of OMPs may have essential functions in Liberibacter growth, making them attractive targets of plant defense mechanisms.

Previously, antibody-based HLB detection has been developed by targeting a Las “OMP” or “OmpA” (35, 36). However, we found that these are actually the same protein that is similar to LasBamA. Peptides derived from another gene previously identified as Las OmpA (CLIBASIA_04260) have been found to reduce Las acquisition by its insect vector (37). However, this protein lacks a transmembrane domain, as predicted by DeepTMHMM, and therefore was not identified as an OMP in our analysis. We hope our systemic analysis of OMPs, especially the β-barrel OMPs, can clarify the nomenclature (**Table S1**).

Our results suggest that the LasOMP1 N-terminal region, including the extracellular O1 loop and the periplasmic P1 domain, mediates both the interaction and cleavage by *Cs*RD21a. It is possible that the O1 loop interacts with *Cs*RD21a in the extracellular fluid in planta and cleaved by this host enzyme as a defense response. This cleavage may disrupt the β-barrel structure of LasOMP1 and expose the periplasmic P1 domain, which can be cleaved by *Cs*RD21a subsequently. Future work should further characterize the cleavage site(s) and clarify how LasOMP1-derived peptide(s) may be released from *Cs*RD21a-dependent cleavage to activate immunity.

OMPs are abundant, cell surface-exposed proteins that contribute to membrane integrity, virulence, and nutrient uptake in Gram-negative bacteria and have been shown to be immunogenic in animal systems (24, 38, 39). In *E. coli*, it is estimated that OMPs cover over half of the OM surface. For example, there are >100,000 copies of *Ec*OmpA alone in the OM (23). A human serine protease, called neutrophil elastase (NE), cleaves *Ec*OmpA, resulting in bacterial membrane instability and growth inhibition (40, 41). As such, the serine protease has direct antimicrobial activities. In this study, LasOMP1 cleavage may also reduce the integrity of Las cell membrane, although this possibility can not be tested due to the inability of growing Las in vitro. In addition to the potential antimicrobial activity, our data also suggests that a PAMP may be released from LasOMP1 cleavage to trigger immune activation. Therefore, animal and plant proteolytic enzymes may target bacterial pathogen OMPs as a conserved strategy but activate defense through different mechanisms.

Similar to previous findings that OMPs in animal pathogens undergo extracellular loop diversification (42), the OMP1 homologs also exhibit sequence diversification at the extracellular loop region. Future research should systematically investigate the impact of this diversification on the functions and defense activation of OMP1. Intriguingly, LcrOMP1 has been found via proteomic analyses in the outer membrane vesicles (OMVs) of Lcr (43). *Ec*OmpA is OMV-associated, and its exogenous application to *E. coli* cells has been shown to reduce NE-mediated killing (44), potentially by sequestering the antimicrobial proteases away from the bacterial cell surface to protect cell integrity. It is intriguing to speculate that Liberibacter may also use a similar strategy to compensate PLCP activities by highly expressing OMP1.

Our data show that overexpression of *Cs*RD21a in citrus enhances HLB tolerance, holding the promise of engineering citrus defense against Las. A naturally occurring mutant of LasOMP1, LasOMP1^Ishi-1^, is also cleaved by *Cs*RD21a suggesting that *Cs*RD21a engineering in citrus may be effective against various Las strains. PLCPs are widespread in plants, representing nine major families (45), thus may contribute to defense against a wide variety of pathogens. For example, C14 may be an exciting target for engineering resistance to Lso in tomato and potato. Yet, unlike *Cs*RD21a, we found that C14 also cleaves *Ec*OmpA in the semi-in vitro cleavage assay, suggesting there is more to be uncovered regarding PLCP specificity against OMPs. Taken together, our results discovered a pathogen substrate of secreted proteases in plant defense, shedding lights on the convergence of plant and animal immune systems on targeting bacterial OMPs, and offering opportunities to engineering defense against the devastating HLB pathogen.

## Materials and Methods

### OMP prediction and analyses

For OMP prediction, bacterial proteomes were downloaded from NCBI (https://www.ncbi.nlm.nih.gov/). PFAM alignments for the membrane β-barrel clan (MBB-CL0193) were downloaded, and hmmbuild was used to build the HMM (25). Next, hmmsearch (http://hmmer.org/) was used to scan each proteome for the presence of OMPs using the HMM. The resulting OMPs were screened using DeepTMHMM (26) and only proteins identified by both programs were subjected to further analyses. MUSCLE v5.1 (46) or MAFFT v7.490 (47) alignments were generated, as indicated, of FASTA amino acid sequences.

Subsequent maximum likelihood model FastTree v2.1.11-generated phylogenies (48, 49) were made and visualized using Geneious Prime (https://www.geneious.com/).

### Yeast-two-hybrid

The yeast strain AH109 (Takara) was maintained in YPDA (Takara) medium prior to transformation. Overnight AH109 liquid cultures were sub-cultured until an OD_600_ between 0.5 and 0.8 was reached. The cells were then washed twice in sterile water followed by pelleting at 1,000 x g for 5 minutes prior to resuspension in transformation buffer, which comprised 50% polyethylene glycol (PEG) 3350, 1M lithium acetate pH 7.0, and 10x Tris-EDTA buffer pH 8.0, at a ratio of 8:1:1. Salmon sperm DNA (Invitrogen) was then added to the transformation mixture with ∼2,000 ng of both bait and prey vectors (Takara), and the cells were incubated at 42°C for 45 minutes. The cells were then pelleted and resuspended in 0.9% NaCl before being plated on YPAD medium. Co-transformed colonies were serially diluted in sterile water after normalizing the OD_600_ as indicated for each experiment and re-plated on SD/-Trp/-Leu (SD-2) and SD/- Trp/-Leu/-His/-Ade (SD-4) (Takara). Primers for pGBKT7 cloning can be found in Table S3.

LcrOMP1a/b/c, LsoOMP1, LamOMP1, and LafOMP1 genes were synthesized as gene fragments (Twist). *Cs*RD21a and *Cs*SAG12-1 in the pGADT7 backbone and LasSDE1 in the pGBKT7 backbone were sourced from Clark*, et al.* (19). LasOMP1 in the pGBKT7 backbone was sourced from Dr. Simon Schwizer.

### Apoplastic fluid extraction

*Cs*RD21a and C14 (as well as their catalytic CHN mutants) were cloned into the binary vector pJK268c (provided by Dr. Jiorgos Kourelis) which contains the RNA silencing suppressor P19 in the backbone (50). Expression was driven by a 2x35S CaMV promoter. *Cs*RD21a and C14 were cloned in their full-length forms without a tag into pJK268c via the pICH41308 Golden Gate level 0 acceptor. Primers can be found in Table S3. These constructs were transformed into *A. tumefaciens* GV3101 competent cells and infiltrated in the leaves of four to six-week-old *N. benthamiana* plants, which were grown in controlled conditions at 25°C with a 16-hour light / 8-hour dark cycle. At three days post-inoculation, the leaves were removed, vacuum infiltrated with sterile water, and centrifuged in the barrel of a 20 mL syringe inserted in a 50 mL Eppendorf tube. The flow-through apoplastic fluid from these leaves was then concentrated 2X using 10 kDa concentrator columns (Thermo) and subjected directly to activity-based protein profiling or the semi-in vitro OMP cleavage assay.

### Activity-based protein profiling

Concentrated apoplastic fluid was collected as mentioned above and was incubated for 30 minutes at room temperature with concentration gradients of E-64 (diluted with DMSO) or just DMSO at room temperature. DCG-04 (MedKoo Biosciences, Inc.) was then added to a final concentration of 500 µM with 5 mM dithiothreitol (DTT) and 50 mM sodium acetate pH 5.0. After one hour incubation at room temperature, protein loading dye was added, and the samples were boiled prior to SDS-PAGE analysis. Western blotting was then performed in which active proteases were detected via chemiluminescence imaging using Streptavidin-HRP (Thermo).

### *E. coli* membrane enrichment of OMPs

Tag-free, outer membrane proteins without their signal peptides were N-terminally fused to the *E. coli* OmpA signal peptide, synthesized as gene fragments (Twist), and cloned into the pOPIN-F6-C vector under the control of the T7 promoter (51). Primers can be found in Table S3. These constructs were transformed into *E. coli* C41 competent cells (52) and grown in liquid auto induction media at 30°C until OD_600_ of 0.5-0.7 was reached, at which point the cultures were moved to 18°C overnight. Pellets were collected via centrifugation at 5,500 rpm (6,561 x g) at 4°C and were resuspended in A1 buffer (200 mM Tris, 200 mM glycine, 20% glycerol, 2M NaCl, and 80 mM imidazole) with cOMPLETE Protease Inhibitor Cocktail tablets (Sigma). The cells were sonicated and centrifuged again at 5,000 rpm (2,990 x g) and 4°C for 30 min. The supernatant was then loaded into PA Ultracrimp tubes (Thermo) and ultracentrifuged at 27,500 rpm (77,077 x g) for 10 minutes. The waxy pellet was then resuspended in A1 buffer with 1% DDM (n-dodecyl-β-D-maltoside) and incubated with rotation at 4°C overnight.

### Semi-in vitro protease cleavage assay

*E. coli* membrane fractions containing OMPs were directly incubated with concentrated apoplastic fluid for three hours at room temperature with final concentrations of 5 mM DTT and 50 mM sodium acetate pH 5.0. Loading dye was added to the samples, which were then boiled. These samples were then run on SDS-PAGE gels and analyzed via Western blotting using custom monospecific OMP1 antibodies raised in rabbits (Pacific Immunology). LasOMP1-specific antibody “⍺-LasOMP1-O3” was raised against the synthetic peptide GPDVAQKYETGKAGEIT, LasOMP1-specific antibody “⍺-LasOMP1-P1” was raised against the synthetic peptide DPVRRAHHGGRGVVPTIATN, and LcrOMP1-specific antibody “⍺- LcrOMP1” was raised against the synthetic peptide RLEYRYTRLGKKDFTLRDA.

### Quenched peptide cleavage assay

Apoplastic fluid containing wild-type or CHN mutant *Cs*RD21a was collected as above and incubated with 4.0 µM E-64 or DMSO for 30 minutes. 5 mM dithiothreitol (DTT) and 50 mM sodium acetate pH 5.0 was then added to 1 µM synthetic peptides QP2 (LKINSAKD) or q-OMP1-O1 (SFGEEAHH). Each peptide was synthesized with an N-terminal DABCYL quencher and a C-terminal Glu-EDANS fluorophore. Fluorescence was immediately measured every minute for 30 minutes at room temperature using a SpectraMax ID5 plate reader with a 335 nm excitation wavelength and a 493 nm emission wavelength. QP2 was sourced from Buscaill*, et al.* (11), and q-OMP1-O1 was generated for this study (Genscript).

### OMP1 detection in healthy and infected citrus

Fruits from infected grapefruit and lemon trees were collected from groves at UF CREC in Lake Alfred, Florida, USA. The seeds were isolated from the fruits, and the seed coats were carefully removed with forceps. The seed vasculatures were then collected with forceps, pooled together, and frozen in liquid nitrogen. These samples were then ground with a mortar and pestle and put on ice. After adding protein loading dye to the samples, they were boiled for 10 minutes and centrifuged at 16,100 x g for 5 minutes. The supernatant was then subjected to SDS-PAGE and subsequent Western blot analyses using the LasOMP1-specific antibodies as above.

### RNAseq analysis of *L. crescens*

Duplicate 3 mL *Liberibacter crescens* BT-1 cultures were grown in liquid BM7 media for 7 days shaking at 150 rpm at 28°C (53, 54). Cultures were spun down at 15,000 x g for 5 minutes at 4°C, and RNA was extracted from bacterial pellets using a Trizol-based extraction protocol (28). The RNA extracts were then treated with DNase I (Thermo Fisher Scientific). The RNA concentration and the quality of the extracts were determined using a NanoDrop spectrophotometer and an Agilent 2100 Bioanalyzer with prokaryote analysis software. RNA libraries were constructed with a TruSeq Stranded Total RNA Library Prep Kit (Illumina). The RNA samples were treated with the Illumina Ribo-Zero Plus rRNA Depletion Kit as part of the standard protocol. cDNA libraries were then subjected to RNA-seq with an Illumina HiSeq system with 150-bp strand-specific paired end reads. Library preparation and sequencing were performed by Genewiz.

The quality of the raw sequencing data was checked with FastQC v.0.11.9 (55). To ensure high-quality sequences for mapping and downstream analyses, low-quality reads and an adapter were trimmed using Trimmomatic v.0.39 (56). RNA-seq reads were aligned with the indexed Lcr BT-1 genome assembly (NC_019907.1) and were parsed to it using HISAT2 v.2.2.1 (57). The number of reads mapped to each gene was counted using the feature counts function of the Rsubread package (58). The data used for the analysis have been deposited into the European Nucleotide Archive database (BioProject PRJEB82095). Read counts were normalized across all samples using the counts function in DESeq2 package v.1.30.1 from Bioconductor (59). Data can be found in Table S2.

### Generation and infection of transgenic citrus

To generate constructs for plant transformation, the full-length coding sequence of *CsRD21a* was amplified from sweet orange with gene-specific primers and cloned into the pDONR221 vector and then fused into the binary vector pK7WG2D using BP and LR enzymes. The *A. tumefaciens* strain EHA105 carrying the recombinant plasmid was used for sweet orange transformation as described previously (60). *CsRD21a*-transgenic citrus plants were confirmed using RT-qPCR with gene-specific primers. Total RNA was extracted from transgenic and WT citrus using Trizol iso plus (Takara). Total RNA was reverse transcribed to cDNA using HiScript II Reverse Transcriptase Kit (Vazyme Biotech). The RT-qPCR was performed with Hieff® qPCR SYBR Green Master Mix (YEASEN) and Light Cycler 480 (Roche). With the citrus β-Actin gene as the internal reference gene, the data were normalized using the 2^−ΔΔCt^ method. All primers used are listed in Table S3. All the transgenic and control plants were grown in a greenhouse at 25-30°C.

### HLB inoculation and Las titer evaluation

For the HLB infection assay, 2-year-old *CsRD21a*-OE and wild-type control plants were grafted as scions on single branches of individual HLB-symptomatic 8-year-old sweet orange trees in the field at the Science Research Institute of Ganzhou, Jiangxi province, China. The buds of transgenic and control lines were grafted at random positions on the same Las-infected branch. After grafting, sampling was initiated once the transgenic and WT buds had developed to a sufficient size, which generally occurred following a four-month growth period. Four months after grafting, leaf midrib samples were collected monthly from both *CsRD21a*-transgenic and WT citrus plants for quantification of Las bacterial titers. In addition, we also grafted HLB-infected citrus buds with similar level of Las titers as scion onto approximately 3-year-old *CsRD21a*-OE and wild-type control plants. Leaf midrib samples were collected monthly from both *CsRD21a*-OE and wild-type citrus plants for quantification of Las bacterial titers. The Las-infected citrus was obtained from an HLB-symptomatic 8-year-old sweet orange tree maintained at the Science Research Institute of Ganzhou, Jiangxi province, China.

DNA was isolated from the leaf midribs monthly after graft inoculation and used to quantify Las titers by Taqman qPCR with primer/probe combination. Las quantification was carried out as follows: DNA was used for qPCR amplification using 16S rRNA primers, the probe HLBp, TaqMan PCR master mix, and SYBR green PCR master mix. The qPCR assays were performed with Light Cycler 480 (Roche) using the SYBR Green PCR Master probe mix (YEASEN) in a 10-µL volume. The data were normalized to the expression of the citrus mitochondrial cytochrome oxidase gene (*COX*). The standard amplification protocol was 95°C for 10 min followed by 40 cycles at 95°C for 10 s and 60°C for 30 s. The bacterial content of Las was represented as CT values, which negatively reflect the bacterial content. The primer details can be found in Table S3.

### *PR1* expression analysis in *N. benthamiana*

The LasOMP1 protein without its signal peptide was fused to the N-terminal PR1 signal peptide and was co-expressed with *Cs*RD21a or C14 in *N. benthamiana.* Primer details can be found in Table S3. GFP and the protease catalytic mutants were used as negative controls. All proteins were expressed using the vector pJK268c in *A. tumefaciens* GV3101 cells, following the same procedure for infiltration in *N. benthamiana* leaves as described above. RNA was extracted from the agroinfiltrated leaves after two days using the RNeasy Plant Mini Kit (Qiagen). Then, 1.0 μg of the extract was digested with 4× gDNA wiper (Vazyme Biotech) to remove gDNA. 5× HiScript II Q RT SuperMix was added to synthesize first-strand complementary DNA (cDNA). The relative expression of *PR1* gene was quantified using RT-qPCR with Taq Pro Universal SYBR qPCR Master Mix (Vazyme) and the CFX Opus 96 Real-Time PCR System (Bio-Rad). RT-qPCR was performed using gene-specific primers (Table S3). Equal amounts of cDNA from nine independent biological replicates were analyzed. Three technical replicates were performed for each biological replicate. With the *N. benthamiana* β-Actin gene as the internal reference gene, relative expression levels were calculated using the 2^−ΔΔCT^ method (61).

### AlphaFold structural modeling

All protein structure modeling was performed using AlphaFold3 (62). Predicted models were imaged using ChimeraX (63–65) and added to the “Gram-negative bacteria cell wall” template image sourced from BioRender.com.

## Acknowledgments

This work is supported by the USDA-NIFA award 2020-70029-33197 to W. Ma and A. Levy. W. Ma is also supported by the Gatsby Charitable Foundation and the UKRI BBSRC grants BB/W00691X/1 and BBS/E/J/000PR9797. N. Wang is supported by the Emergency Citrus Disease Research and Extension program, project award no. 2022-70029-38471 and 2023-70029-41280 from USDA-NIFA, Florida Citrus Initiative Program, Citrus Research and Development Foundation, and Hatch project (FLA-CRC-005979). R. van der Hoorn is supported by ERC project 101019324. Q. Xu is supported by the National Natural Science Foundation of China (32525008 and U23A20198). We thank Drs. Jiorgos Kourelis, Simon Schwizer, Adam Bentham, Mauricio Contreras, and Yinghong Gu for experimental guidance and support, the TSL support teams and JIC Horticultural Services for plant growth, and Amanda Chai and Priya Desikan for experimental assistance. We are indebted to past and present members of the Ma, Levy, Wang, Xu, van der Hoorn, and Hogenhout labs for their support, advice, and discussions.

## Author Contributions

AJM, BH, YX, QX, and WM designed the research; AJM, BH, YX, CW, EH, YW, and ZP performed the research; RVDH, AL, and NW contributed to new reagents/analytic tools; AJM, BH, YX, AHL, and ADF analyzed data; AJM, BH, YX, QX, and WM wrote the paper.

---

**Figure S1.**
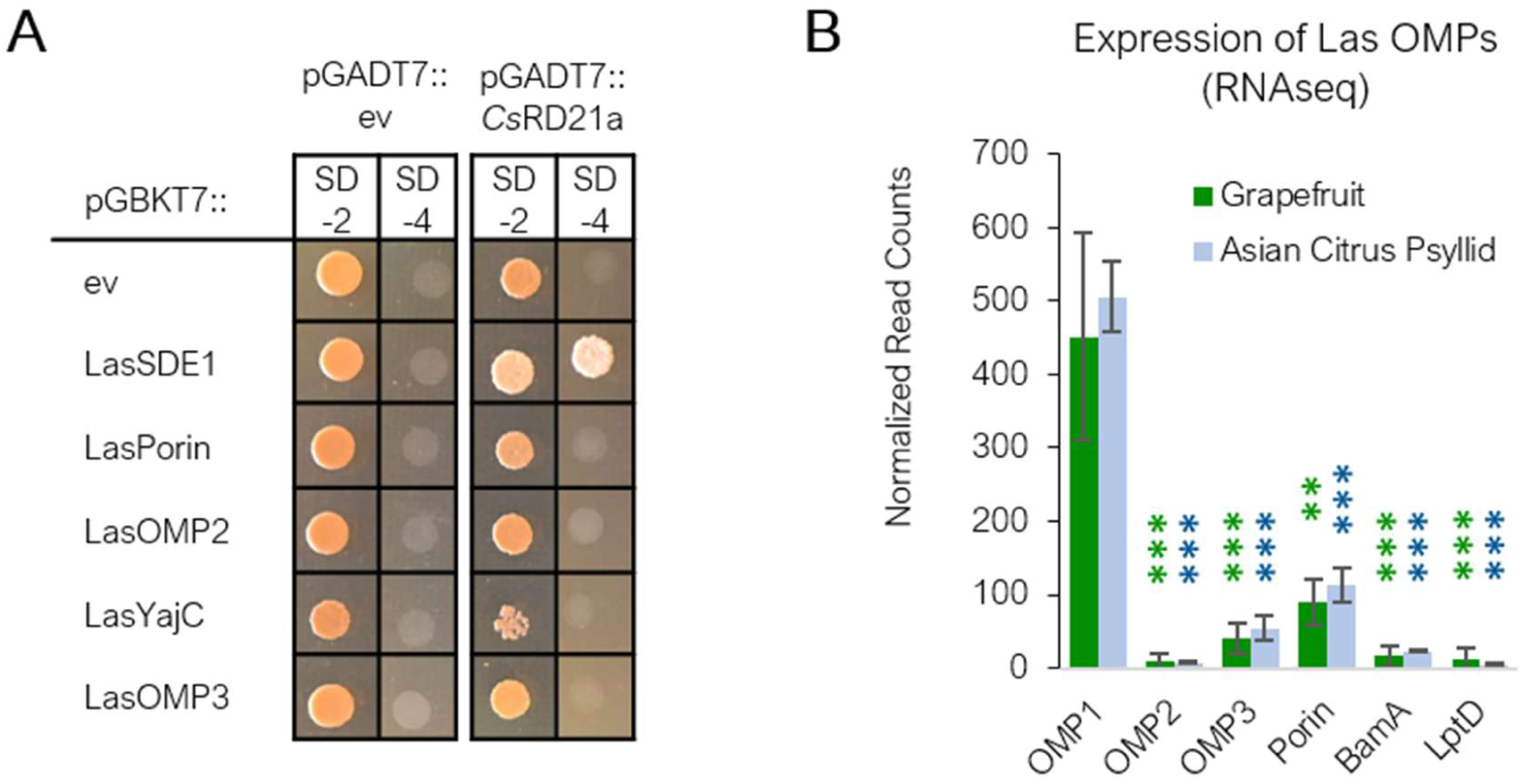
*Cs*RD21a does not interact with other Las OMPs, which exhibit lower expression than LasOMP1. (**A**) Yeast-two-hybrid assay examining interaction of *Cs*RD21a (prey) with Las OMPs (bait) with LasSDE1 as the positive control and empty vector (ev) as the negative control. LasYajC, an inner membrane protein, was also examined. SD-2 medium lacks leucine and tryptophan and was used to select the co-transformed colonies. One colony with OD_600_ = 1.0 was plated for each co-transformation in yeast on both SD-2 and SD-4, which lacks leucine, tryptophan, adenine, and histidine. All proteins were expressed without their signal peptides. (**B**) Expression of Las OMPs in infected grapefruit midribs (n=6) and the Asian citrus psyllid (n=4) represented by normalized read counts using publicly available data (27). Error bars indicate standard deviation. Asterisks (*) represent degrees of significance as determined by two-tailed T-test between each gene and LasOMP1, where ** signifies p < 0.01 and *** signifies p < 0.001. Green and blue asterisks represent significance for the grapefruit and Asian citrus psyllid samples, respectively.

**Figure S2.**
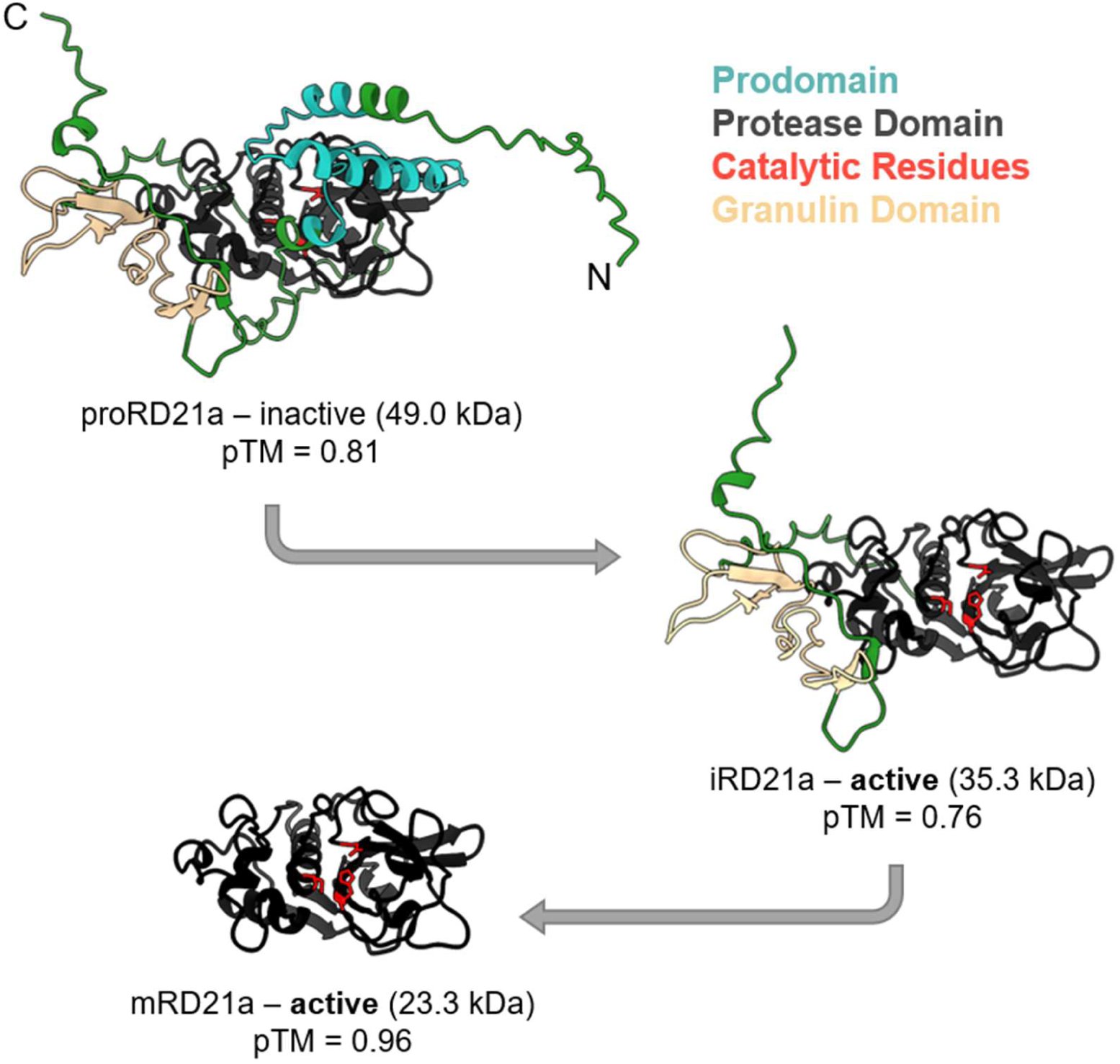
Processing of *Cs*RD21a in planta. *Cs*RD21a *s*tructures were predicted with AlphaFold3 and visualized using ChimeraX. RD21a proteins have an N-terminal signal peptide (not shown), an autoinhibitory prodomain (blue), a catalytic protease domain (black), three catalytic residues (red), and a C-terminal granulin domain (tan). proRD21a (shown at the top) is generated when the N-terminal signal peptide has been removed, but the enzyme is inactive due to the presence of the autoinhibitory prodomain. iRD21a is an active intermediate in which the prodomain has been removed, exposing the catalytic residues. mRD21a is a further processed, active enzyme that has had its granulin domain removed. The protein molecular weights (in kDa) and AlphaFold3 confidence scores (in pTM) are shown. N- and C-termini are labeled on proRD21a.

**Figure S3.**
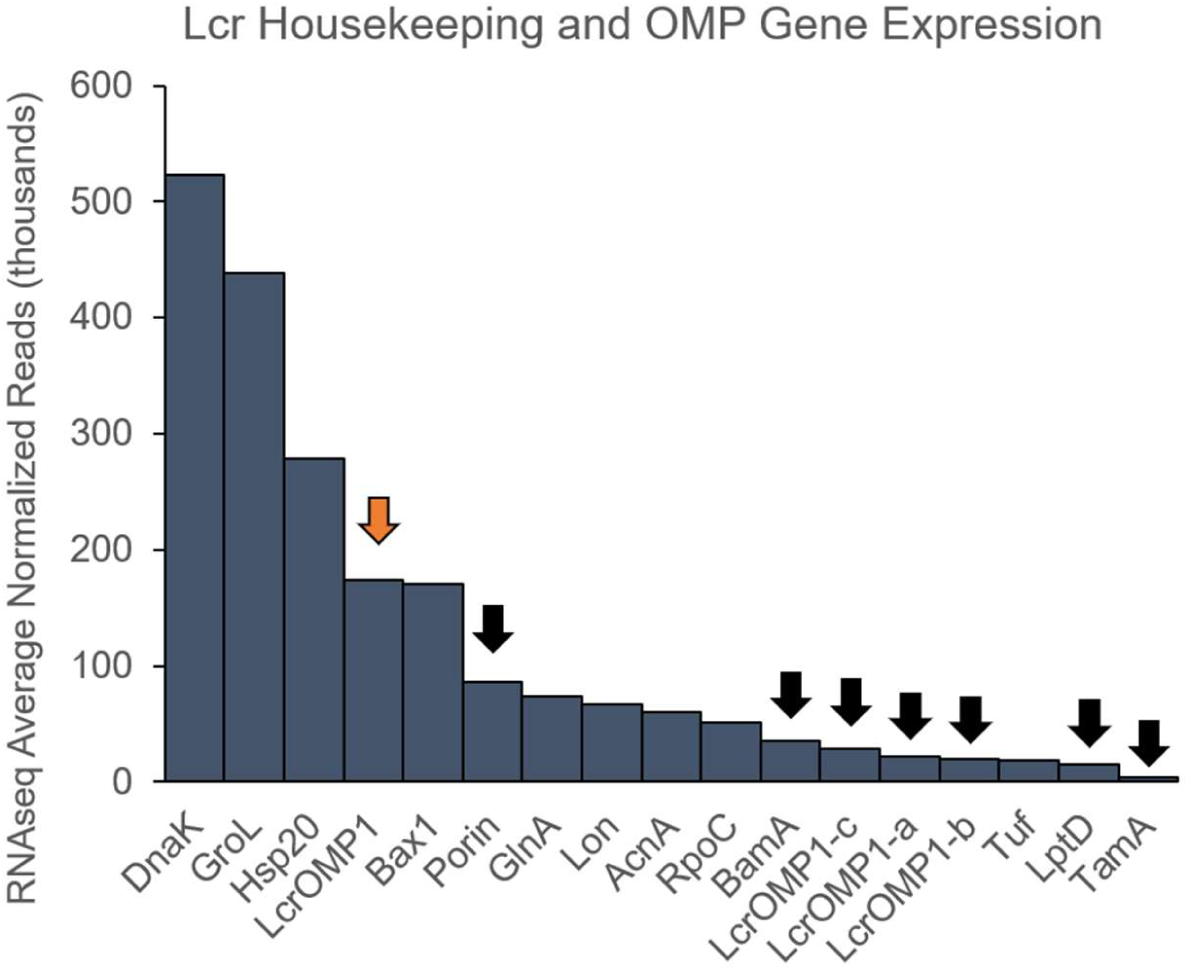
LcrOMP1 is highly expressed. Expression levels, represented by average normalized reads (two replicates), of genes encoding OMPs (marked with black arrows) in Lcr grown in artificial medium as determined by RNA-seq. Some of the most highly expressed housekeeping genes are used as a comparison. All OMPs, except for TamA, are in the top 17% of highest expressed genes in the dataset (Table S2).

**Figure S4.**
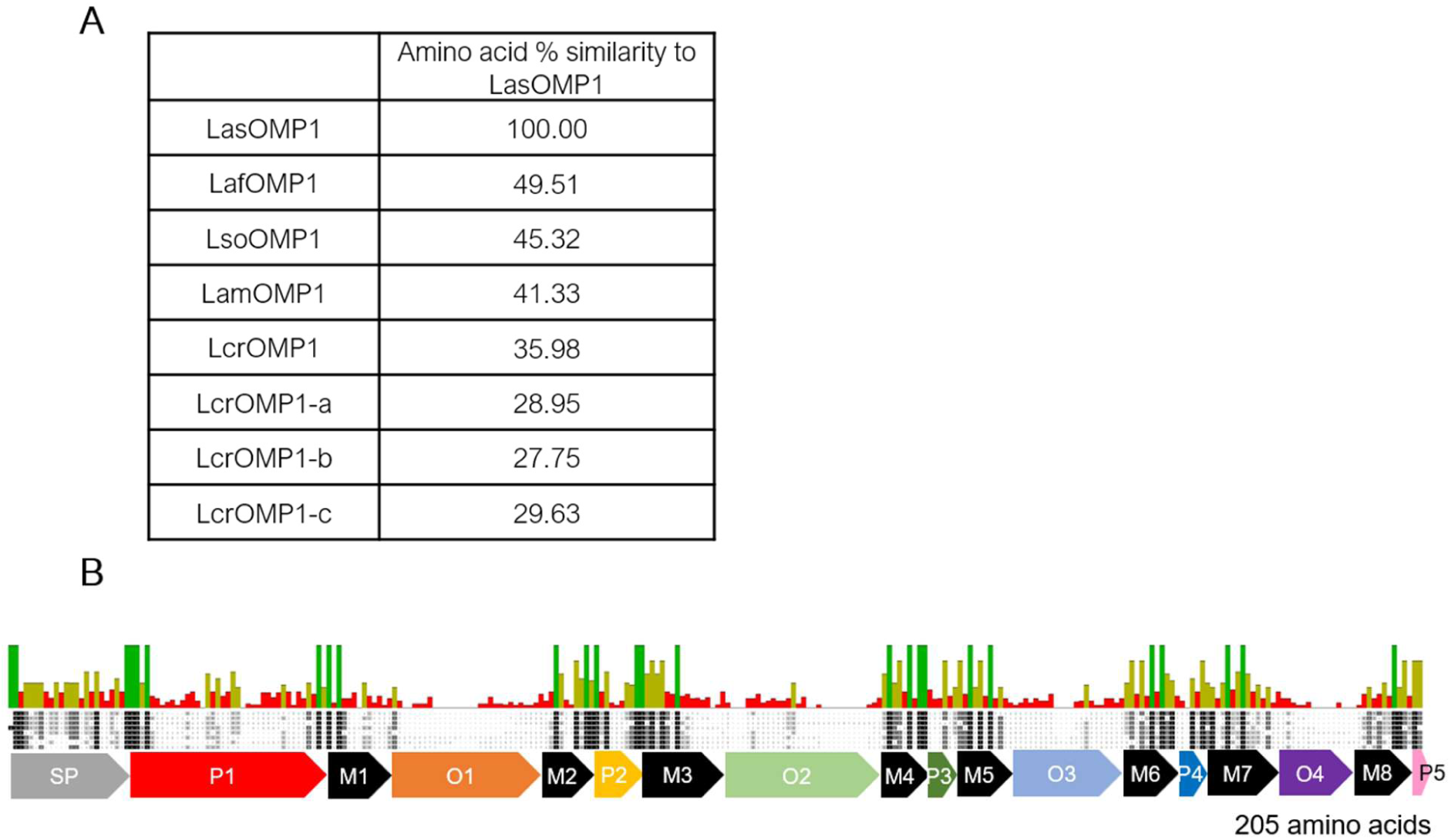
Sequence comparison of OMP1 homologs in *Liberibacter* spp. (**A**) Amino acid sequence similarity as determined by MUSCLE alignment (46) of OMP1 homologs. (**B**) MUSCLE alignment of OMP1 homologs mapped to the domain structure of LasOMP1. Conservation levels are displayed for each amino acid (green = 100% conserved, green-brown = 30-99% conserved, red = 0-29% conserved).

**Figure S5.**
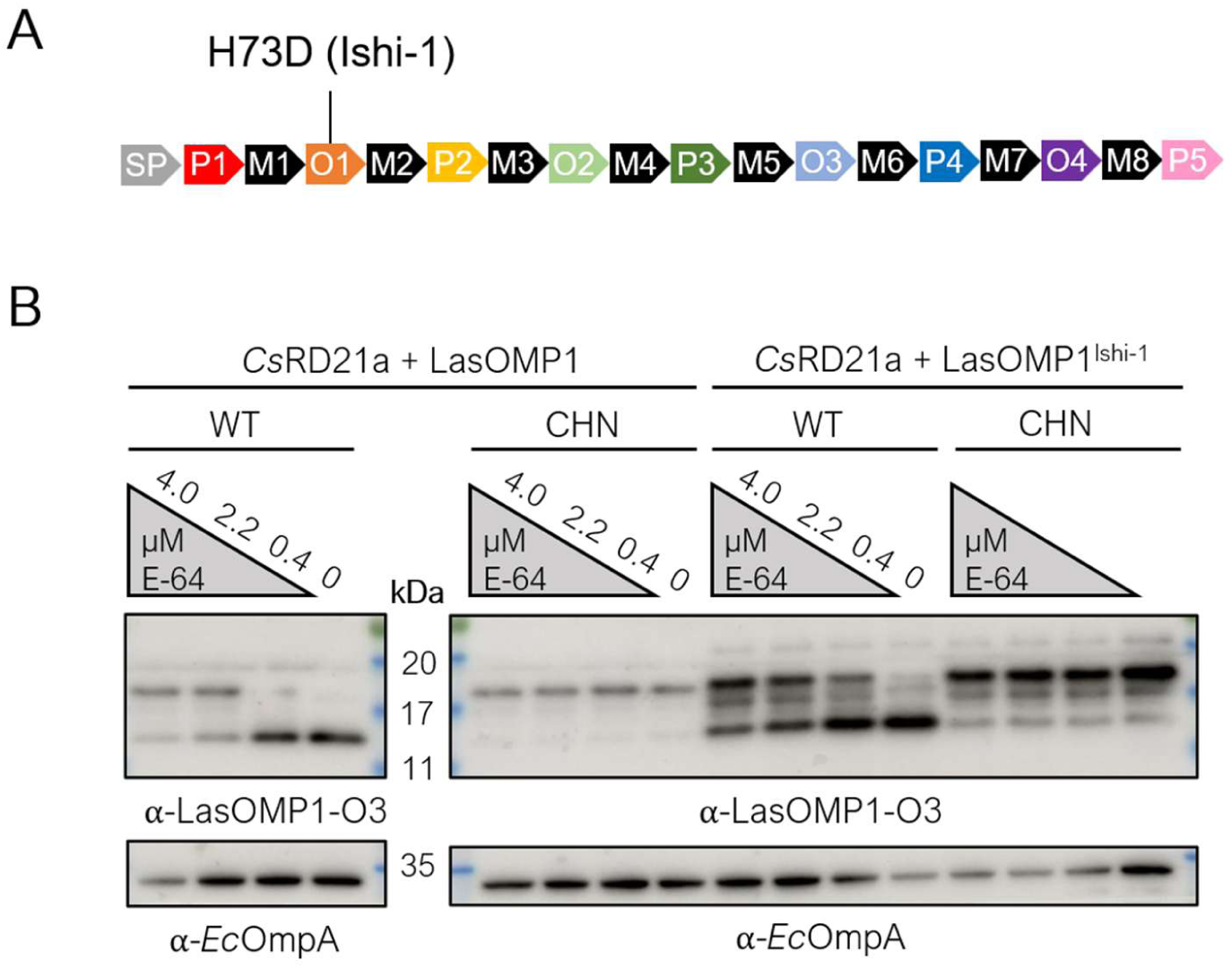
*Cs*RD21a cleaves LasOMP1^Ishi-1^. (**A**) Domain structure of LasOMP1 (strain Ishi-1) depicting a non-synonymous mutation found in the O1 domain. (**B**) Semi*-*in vitro protease cleavage assay of *Cs*RD21a with LasOMP1 or the LasOMP1^Ishi-1^ variant. E-64 or DMSO-treated apoplastic fluid containing *Cs*RD21a wild-type (WT) or CHN mutant (CHN) was incubated with LasOMP1 or the LasOMP1^Ishi-1^ variant from *E. coli* membrane fractions, and *Ec*OmpA was used as a loading control. Both OMPs were detected via Western blot using protein-specific antibodies.

**Figure S6.**
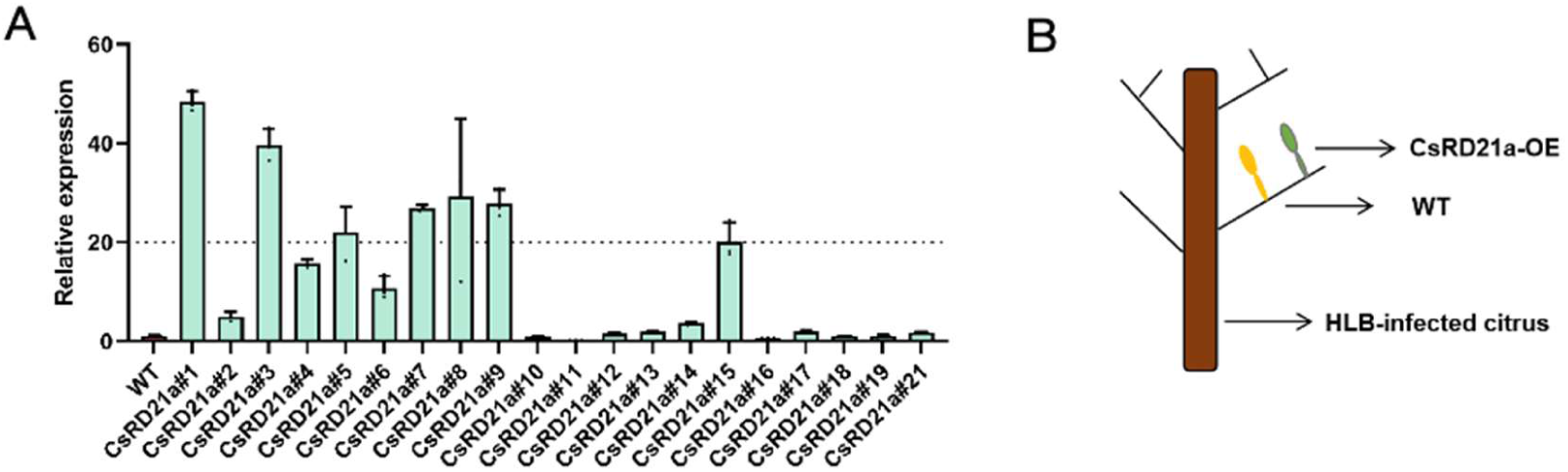
Confirmation of *CsRD21a* overexpression in transgenic sweet orange plants. (**A**) Relative expression of *CsRD21a* in 20 independent *CsRD21a*-overexpressing (OE) lines compared to wild-type (WT) plants. Expression was determined by RT-qPCR and normalized to the citrus *β-Actin* gene. Data represent means ± standard error (SE) from three biological replicates. Lines 1, 3, 5, 7, 8, and 15 were used for subsequent Las infection assays. (**B**) Schematic diagram showing the grafting of *Cs*RD21a-OE or WT scions on the same branch of HLB-infected citrus.

**Figure S7.**
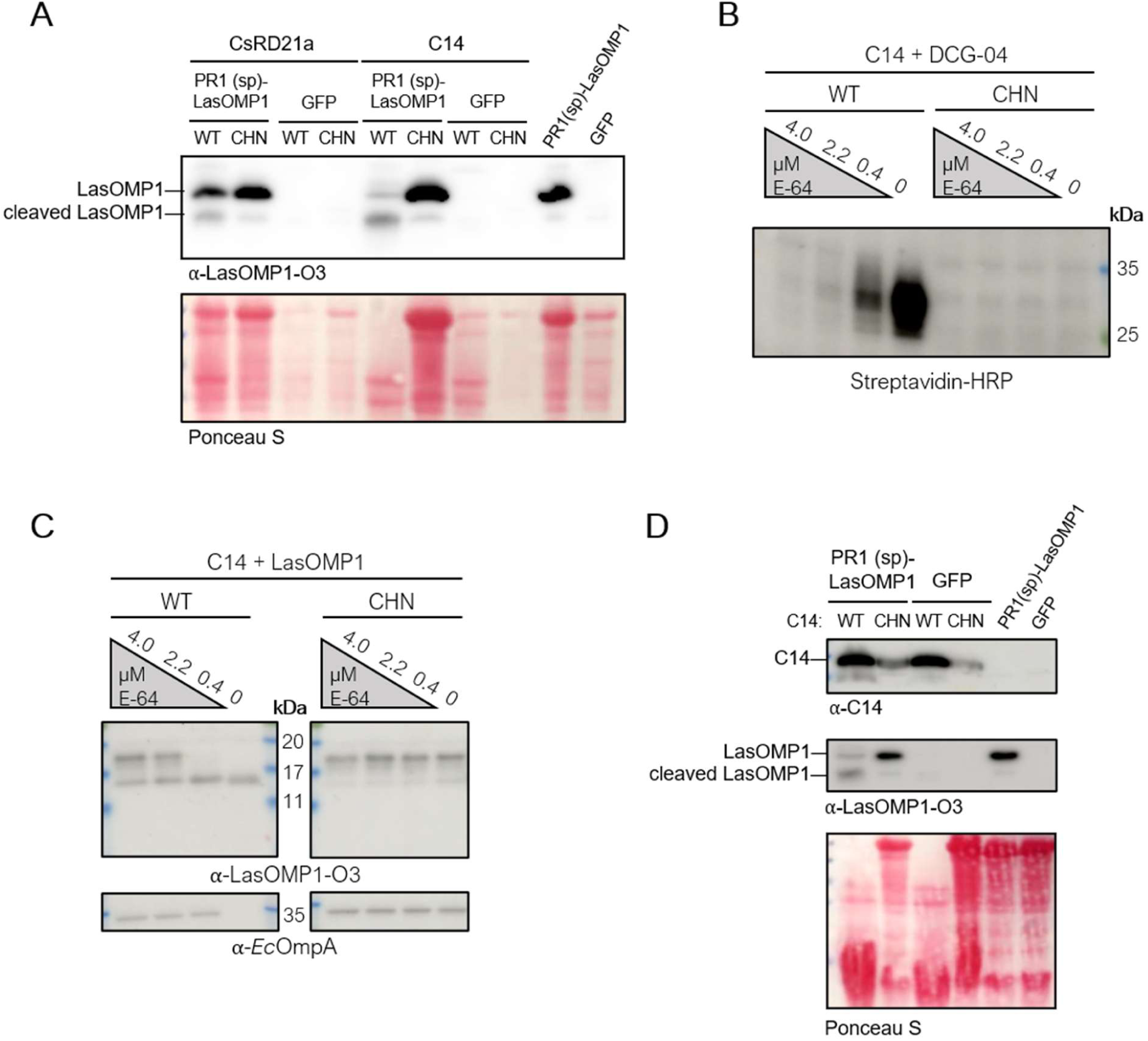
Tomato RD21a (C14) cleaves LasOMP1. (**A**) Western blot depicting LasOMP1 cleavage by *Cs*RD21a or C14. LasOMP1 fused to the signal peptide (SP) from PR1, instead of its native SP, was co-expressed with wild-type (WT) or catalytic mutant (CHN) *Cs*RD21a or C14 in *N. benthamiana*. Green fluorescent protein (GFP) was used as a negative control. (**B**) Western blot detection of wild-type (WT) or catalytically dead (CHN) C14 in apoplastic fluid (AF) using activity-based protein profiling (ABPP), where a concentration gradient of the cysteine protease inhibitor E-64 was incubated with the AF prior to DCG-04 labeling. Western blot detection of DCG-04-bound proteases was performed using Streptavidin-HRP. (**C**) LasOMP1 is cleaved by C14 in the semi-in vitro assay. LasOMP1 was expressed in *E. coli* and the resulting membrane fractions were incubated with E-64- or DMSO-treated, C14-containing apoplastic fluid. *Ec*OmpA could also be cleaved by C14, but only in the absence of E-64. (**D**) Confirmation of C14 expression when the WT protein or CHN mutant is co-expressed with LasOMP1 or GFP in *N. benthamiana*. These samples were included in the RT-qPCR analysis in Fig. 5E.

**Table S1. HMM-predicted OMPs in Liberibacter.** Beta barrel OMPs predicted by both hmmsearch and DeepTMHMM in representative Liberibacter strains: CLas (strain psy62), CLaf (strain PTSAPSY), CLso (strain ISR100), CLam (strain PW_SP), and Lcr (strain BT-1). Annotations are derived from NCBI reference genomes for each strain, and clusters are colored based on Figure 3A.

**Table S2. Normalized counts of 1,388 genes in *Liberibacter crescens* BT-1 transcriptome for duplicate bacterial samples collected from liquid culture.** Genes are ordered based on average normalized counts in descending order. The normalized counts were calculated using counts function in DESeq2 v.1.30.1 R package. Gene, gene names, old locus names, and description were taken from NCBI.

**Table S3. Primers used in this study.**

